# TRIM52 downregulates IFN-β Production by Targeting TBK1 for Proteasome Degradation

**DOI:** 10.64898/2026.06.24.734385

**Authors:** Qian Qin, Chunfu Zheng

**Affiliations:** College of Medicine, Lishui University, Lishui, 323000, China; Department of Microbiology, Immunology, and Infectious Diseases, University of Calgary, Calgary, Alberta, Canada

**Keywords:** TRIM52, TBK1, ubiquitination, degradation, proteasome

## Abstract

The IFN-I (type I interferon) signaling pathway is the first line of defence against foreign pathogens. Stringent control of signalling pathways is necessary to maintain host immune responses and homeostasis. However, the underlying mechanism for its tight regulation is yet completely understood. In this study, we demonstrated that the TRIM family protein tripartite motif-containing 52 (TRIM52) is a novel negative regulator of IFN-β production. Ectopically expressed TRIM52 markedly inhibited the activation of the IFN-β promoter by ectopic expression of cGAS/STING, RIG-IN, or TRIF, MAVS, STING, and TBK1 but not by IRF3/5D, indicating that TRIM52 targets TBK1. TRIM52 also significantly inhibited the IFN-β, ISG54, and ISG56 production, the dimerization of IRF3 and the nuclear localization of IRF3-YFP induced by ectopic expression of TBK1. Co-immunoprecipitation experiment revealed that TRIM52 specifically interacted with TBK1. Furthermore, the TBK1 protein, but not its mRNA, decreased considerably with increasing expression of TRIM52, and TRIM52 did not decrease the expression of the cGAS, STING, or IRF3 proteins. In addition, proteasome inhibitor MG-132 blocked the reduced TBK1 induced by TRIM52, indicating that TRIM52 caused TBK1 degradation via the proteasome pathway. Co-IP and ubiquitination assays demonstrated that TRIM52 promotion of K48-linked ubiquitination of TBK1, which depends on its E3 ubiquitin ligase. Collectively, our findings identify a previously unrecognized role of TRIM52 in regulating the IFN-I signalling pathway through targeting TBK1 for polyubiquitination and degradation.

## Introduction

The innate immune system is the first line of host defence against invading pathogens. Initiation of the innate immune response depends on pattern recognition receptors (PRRs), including Toll-like receptors (TLRs), RIG-I-like receptors (RLRs), Nod-like receptors (NLRs) and DNA sensors, to recognize pathogen-associated molecular patterns (PAMPs) [1, 2]. Upon recognizing PAMPs, PRRs activate downstream IFN-I, transcription factor NF-κB and inflammasome signalling pathways, leading to the production of IFN-stimulated genes (ISGs) and proinflammatory cytokines and inducing an antiviral immune response. TLR3 recognizes dsRNA to activate the downstream adaptor protein Toll/IL-1 receptor domain-containing adaptor-inducing IFN-β (TRIF); RIG-I and MDA5 recognize cytoplasmic dsRNA to activate the mitochondrial signalling adaptor MAVS; and several viral DNA sensors, including DEAD-box helicases (DDX41), IFN-γ-inducible protein 16 (IFI16) and Cyclic GMP-AMP synthase (cGAS), recognize cytosolic DNA to activate the membrane-associated adaptor stimulator of interferon genes (STING). All of the key adaptors TRIF, MAVS, and STING recognize the kinase TBK1; subsequently, the activation of the transcription factors interferon regulatory factor 3/7 (IRF3/7) and NF-κB leads to the production of IFN-I and inflammatory cytokines [3, 4].

Multiple post-translational modifications, especially ubiquitination, have been reported to regulate the activity of TBK1. NLRP4/DTX4 or TRIP mediate K48-linked ubiquitination of TBK1 [5, 6], resulting in the degradation of TBK1 and inhibition of the TBK1-IRF3 signalling cascade. On the other hand, TBK1 undergoes Lys (K) 63 linkage ubiquitination induced by Ndrp1 and mind bomb in response to TLR and RLR ligands, respectively, and promotes IFN-I production [7, 8]. Although studies have reported on the ubiquitination of TBK1, more details regarding TBK1 ubiquitination need to be elucidated to understand the regulation of innate immune responses better.

Tripartite interaction motif (TRIM) proteins are RING-type E3 ligases that share an N-terminal structure consisting of a RING domain, one or two B boxes, a putative coiled-coil domain, and a variable C-terminus; more than 70 TRIM proteins have been identified in humans [9]. TRIM proteins are involved in diverse cellular processes, including autophagy, antiviral activity, autoimmune diseases, and oncogenesis [10]. Accumulating evidence suggests that many TRIM proteins are involved in the immune system. In addition, several TRIM proteins have been reported to modulate TBK1 activity in different ways. Upon viral infection, TRIM9s undergo Lys63-linked autopolyubiquitination and serves as a platform to bridge GSK3β to TBK1, leading to the activation of IRF3 signalling. TRIM11 interacts with and inhibits the activity of TBK1-adaptor complexes, reducing the IFN-induced antiviral state against HSV-1 and VSV-GFP [11, 12], and TRIM26 reportedly also interacts with TBK1 and restricts RNA virus infection by binding to TBK1 and NEMO to facilitate downstream signalling [13]. However, whether TBK1 is regulated by other TRIM proteins and the detailed mechanisms of action remain largely unknown [14, 15].

TRIM52 contains the RING domain and B-box domain [16], and TRIM52 mRNA is expressed at readily detectable, moderately low levels in all tested cell lines as the protein is rapidly turned over by the proteasome [17]. To date, TRIM52 has been shown to promote oncogenesis, inhibit viruses and upregulate NF-κB activation [18-20]. In this study, we demonstrated that TRIM52 interacts with TBK1 and targets it for K48-linked ubiquitination and degradation, downregulating IFN-I signalling in response to both DNA and RNA viruses as TBK1 are share by both DNA and RNA-sensing signalling pathways.

## MATERIALS AND METHODS

### Cells and antibodies

HEK293T was obtained from the American Type Culture Collection (Manassas, VA) and cultured in Dulbecco’s modified Eagle medium (DMEM) (Gibco-BRL) supplemented with 10% fetal bovine serum (FBS) and 100 U/ml penicillin and streptomycin. The proteasome inhibitor MG132 was purchased from Thermo Fisher Scientific (USA). Radio immunoprecipitation assay (RIPA) lysis buffer was purchased from Beyotime (Shanghai, China). Mouse anti-Myc, anti-Flag, anti-HA, and anti-GAPDH monoclonal antibodies (MAbs) were purchased from Abmart (Shanghai, China). Rabbit anti-IRF3 and anti-TBK1 polyclonal antibodies (PAbs) were obtained from GL Biochem Ltd. (Shanghai, China).

### Plasmid construction

TRIM52-FLAG/MYC expression plasmids were constructed by standard molecular biology techniques. The commercial reporter plasmid pRL-TK (RL stands for Renilla luciferase, and TK stands for thymidine kinase) was purchased from Promega Corp. The other plasmids used included the following: Rongtuan Lin provided the FLAG-TBK1, IRF3/5D-FLAG, and IFN-β luciferase reporter plasmids as gifts.

### Co-IP and Western blot analysis

Co-IP assays and WB analysis were performed as previously described [27]. In brief, HEK293T cells were seeded and cotransfected with indicated plasmids encoding indicated proteins, and the cells were lysed on ice in lysis buffer (1.5 M Tris [pH 7.4], 1 M NaCl, 0.5 M EDTA [pH 8.0], 6.0 M sodium deoxycholate, and 1 % Triton X-100). Protein A/G PlusAgarose (Santa Cruz Biotechnology) and the corresponding antibodies were then added to the lysates, which were subsequently incubated overnight at 4°C. After three washes with lysis buffer, the beads were subjected to the WB analysis.

### RNA isolation and qRT-PCR

Total RNA was extracted using TRIzol (Invitrogen) according to the manufacturer’s instructions. The samples were dissolved in RNase-free water, digested with DNase I, and then subjected to reverse transcription as previously described. cDNA was used as the template for qRT-PCR to measure the levels of IFN-β and ISGs, and 18S rRNA was used as an internal reference as previously described. The primers used for qPCR were as follows: IFN-β, forward, 5‘-CCAACAAGTGTCTCCTCCAAAT-3’ and reverse, 5‘-AATCTCCTCAGGGATGTCAAAGT-3’; and ISG54, forward, 5‘-CTGCAACCATGAGTGAGAA-3’ and reverse, 5‘-CCTTTGAGGTGCTTTAGATAG-3’; and ISG56, forward, 5‘TACAGCAACCATGAGTACAA-3’, and reverse, 5‘-TCAGGTGTTTCACATAGGC-3’; TBK1, forward, 5‘-CAACCTGGAAGCGGCAGAGTTA-3’; and reverse, 5‘-ACCTGGAGATAATCTGCTGTCGA-3’; GAPDH, forward, 5‘-GACACCCACTCCTCCACCTTT-3’; and reverse, 5‘-ACCACCCTGTTGCTGTAGCC-3’.

### Transfection and DLR assays

Cells were transfected with Lipofectamine 2000 (Invitrogen) according to the manufacturer’s recommendations. HEK293T cells were transfected with reporter plasmids, such as IFN-β-Luc and the internal control plasmid pRL-TK, with or without expression plasmids, as indicated, by Lipofectamine 2000. At 24 h posttransfection, luciferase assays were performed with a dual-specific luciferase assay kit (Promega, Madison, WI) as described in our previous studies.

### Immunofluorescence assays

Immunofluorescence assays were performed as described previously [28]. In brief, HEK293T cells were transfected with TRIM52-MYC, TBK1-FLAG and IRF3-TFP for 20 h and then rinsed three times with PBS. The cells were then washed with PBS after being fixed in 4 % paraformaldehyde. Primary antibodies were added after the samples had been incubated for 1 h at 37°C to prevent nonspecific binding, after which they were incubated for 8 h at 4°C. After that, the samples were stained with DAPI for ten minutes in the dark at room temperature. Fluorescence microscopy (Zeiss) was used to obtain images.

### Statistical analysis

Data are represented as the means ± standard deviations (SDs) when indicated, and Student’s *t*-test was used for all statistical analyses with GraphPad Prism 5.0 software. Differences between groups were considered significant when the *P*-value was <0.05.

## RESULTS

### TRIM52 downregulates both the DNA and RNA sensing signalling pathways

To identify the possible role of TRIM52 in innate immunity, we examined the effect of the expression of TRIM52 on IFN-β expression using an IFN-β promoter activity assay. HEK293T human embryonic kidney cells (293T cells) were cotransfected with IFN-β luciferase (IFN-β-Luc) reporter, Renilla luciferase reporter (RL-TK), and TRIM52 expression plasmids, along with cGAS/STING, RIG-IN (a RIG-I-activating mutant that has only the RIG-I N-terminal CARD domain), or TRIF for 20 h. We found that ectopic expression of cGAS/STING, RIG-IN, or TRIF activated the IFN-β promoter in HEK293T cells and that cotransfection of TRIM52 markedly inhibited the activation of the IFN-β promoter (Fig. 1A-C). These results indicated that TRIM52 downregulates both DNA and RNA-sensing signalling pathways.

**Fig. 1.**
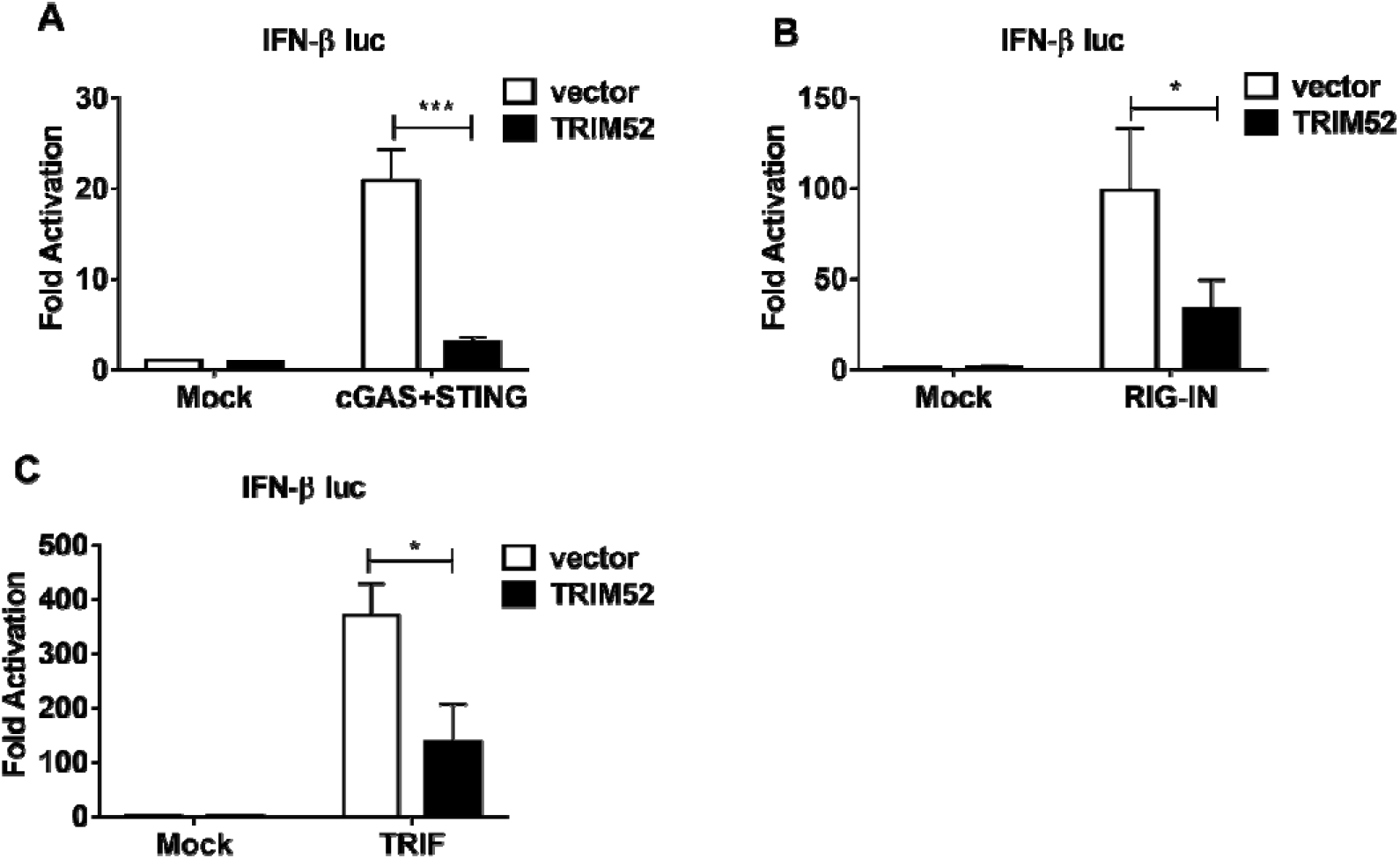
TRIM52 downregulates both the DNA and RNA sensing signalling pathways.

### TRIM52 interacts with and targets TBK1

To determine the molecular mechanisms through which TRIM52 downregulates both DNA and RNA-sensing signalling pathways, we cotransfected HEK293T cells with expression vectors encoding MAVS, STING, TBK1, or IRF3/5D (a constitutively active mutant of IRF3) together with IFN-β-Luc reporter and RL-TK reporter. We found that TRIM52 inhibited IFN-β reporter activity induced by MAVS, STING, and TBK1 but not by IRF3/5D (Fig. 2A-D), indicating that TRIM52 targets TBK1. Quantitative reverse transcription-PCR (qRT-PCR) revealed that TRIM52 significantly inhibited the IFN-β, ISG54, and ISG56 transcription induced by TBK1 (Fig. 2E-G). These results suggested that TRIM52 may inhibit both DNA and RNA-sensing signalling pathways by targeting TBK1.

**Fig. 2.**
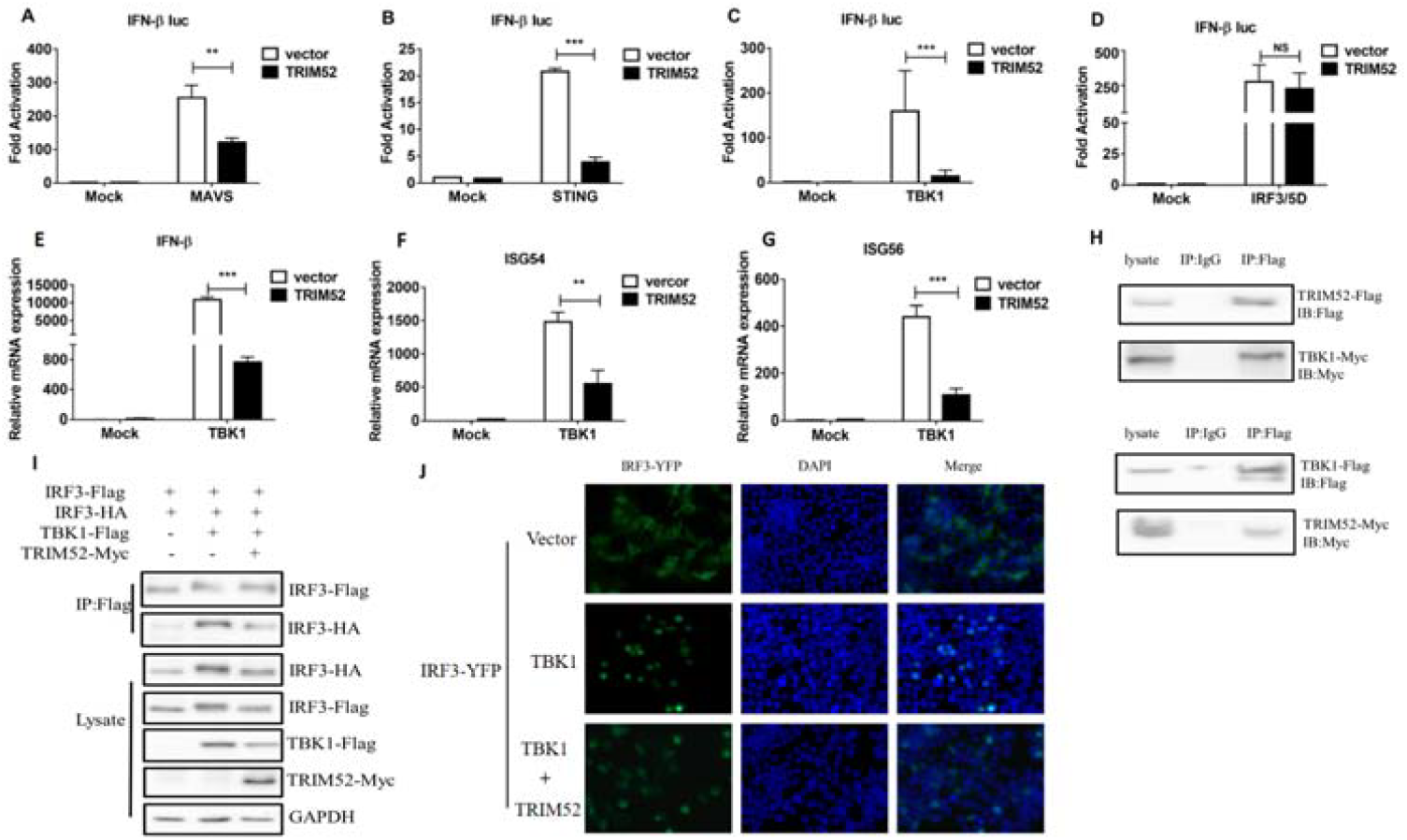
TRIM52 interacts with and targets TBK1.

Next, we sought to determine whether TRIM52 could interact with TBK1, and co-immunoprecipitation experiments revealed that TRIM52 specifically interacted with TBK1 (Fig. 2H). In addition, we assessed the dimerization of IRF3 in HEK293T cells expressing TRIM52 together with TBK1 and found that TRIM52 potently inhibited the dimerization of IRF3 induced by TBK1 (Fig. 2I). Next, an immunofluorescence assay was used to examine the translocation of IRF3-YFP in cells with or without the ectopic expression of TRIM52. As shown in Fig. 2J, in cells transfected with TBK1, IRF3-YFP rapidly translocated from the cytoplasm to the nucleus; in contrast, IRF3-YFP was retained in the cytoplasm of cells with ectopically expressing TRIM52 (Fig. 2J). Taken together, these results indicated that TRIM52 inhibits the activation of IFN-I by targeting TBK1.

### TRIM52 targets TBK1 for proteasome pathway degradation

As TRIM52 negatively regulates both DNA and RNA sensing signalling pathways through targeting TBK1, we examined whether TRIM52 regulates TBK1 protein. HEK293T cells were transfected with a plasmid encoding TBK1 together with increasing amounts of a plasmid encoding TRIM52. As a result, the concentration of the TBK1 protein decreased considerably with increasing expression of TRIM52 (Fig. 3A). To exclude the possibility that the decrease of TBK1 protein was resulted from decreased transcription of the TBK1 gene, we used RT-PCR to analyze the same HEK293T cells expressing various genes and found that the abundance of TBK1 mRNA did not change with increasing expression of TRIM52 (Fig. 3A). In addition, increasing TRIM52 expression did not decrease the transcription of the cGAS, STING, or IRF3 proteins (Fig. 3B-D).

**Fig. 3.**
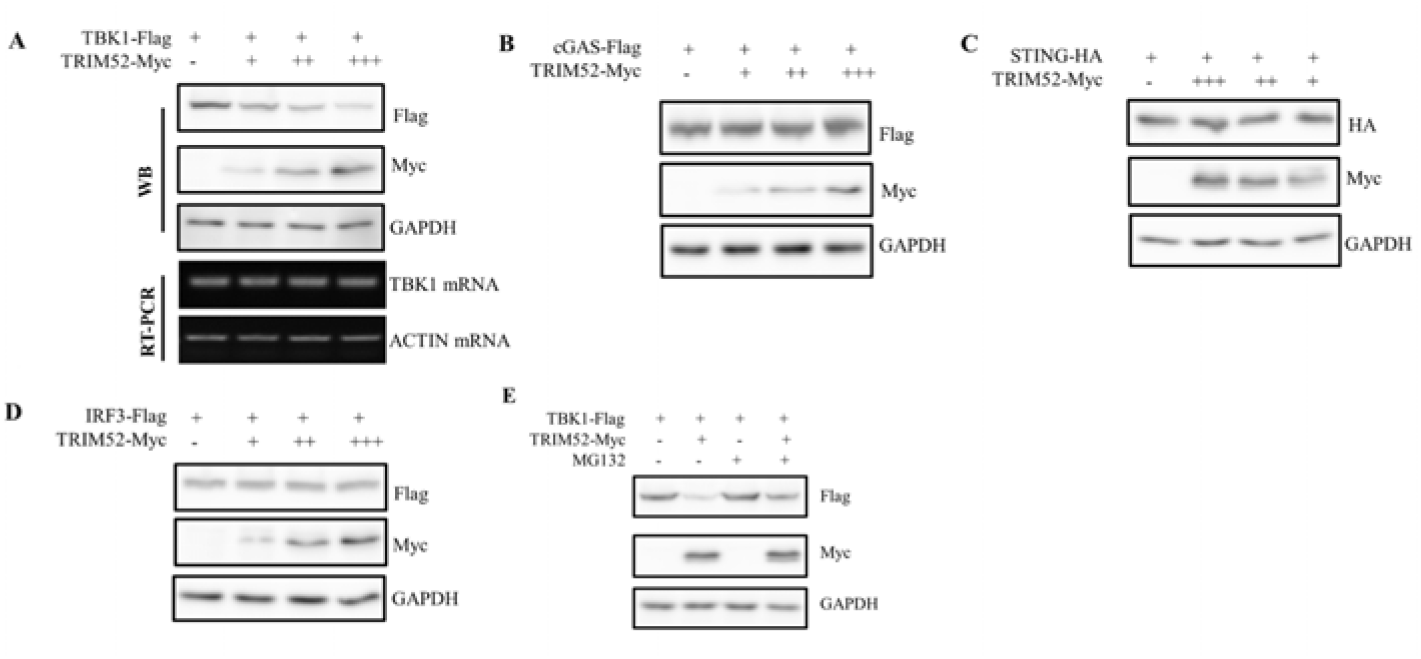
TRIM52 targets TBK1 for proteasome pathway degradation.

The ubiquitination of TRIM52 is critical for its regulatory function [19]. We investigated whether TRIM52-mediated TBK1 regulation is dependent on the ubiquitination of TRIM52. We found that the loss of the TBK1 protein induced by TRIM52 was blocked by the proteasome inhibitor MG-132, indicating that TRIM52 degradates TBK1 via the proteasome pathway (Fig. 3E).

### TRIM52 promotion of K48-linked ubiquitination of TBK1 depends on its E3 ubiquitin ligase

Our above results revealed that TRIM52 degrades TBK1 through the proteasome pathway. To investigate whether the E3 ligase activity of TRIM52 was involved in the regulation of TBK1 activation, we transfected Myc-TBK1 into HEK293T cells together with the FLAG-TRIM52 construct. Co-immunoprecipitation experiments revealed that TBK1 ubiquitination was markedly increased in the presence of the ectopic expression of TRIM52 (Fig. 4A), even with the decreased TBK1. To investigate the underlying mechanism of TRIM52-mediated TBK1 polyubiquitination, we used mutants in which all the lysine residues except K48 were replaced with arginine (HA-ubiquitin-K48) and only the lysine residue K48 was replaced with arginine (HA-ubiquitin-K48R). We detected increased K48-linked ubiquitination of TBK1 in cells co-expressing TRIM52 and TBK1, whereas the amount of K48R-linked ubiquitination of TBK1 remained unchanged compared with that in cells transfected with the TBK1 expression plasmid alone (Fig. 4B).

**Fig. 4.**
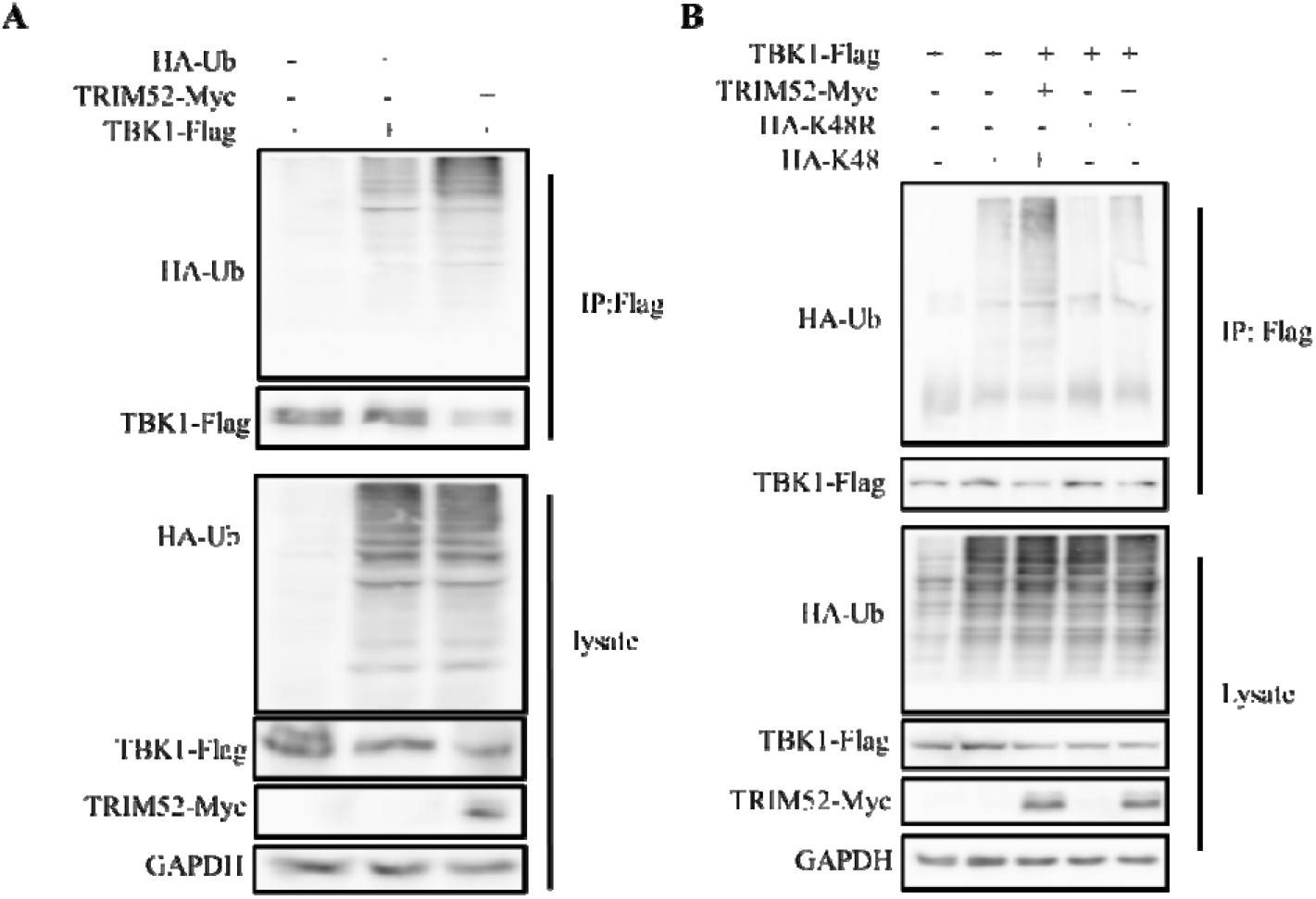
TRIM52 promotes K48-linked ubiquitination of TBK1.

TRIM52 is a RING-type E3 ligase that contains a RING domain and one B box domain. To identify the molecular mechanisms by which TRIM52 targets TBK1 for K48-linked ubiquitination, we generated a TRIM52 deletion mutant without the RING domain (ΔRING domain) and assessed its ability to inhibit TBK1-induced signalling. We found that full-length TRIM52 and RING domain of TRIM52 alone inhibited the TBK1-induced activity of the IFN-β luciferase reporter, but RING domain deletion mutant TRIM52 did not (Fig. 5A). We further found that similar to full-length TRIM52 and RING domain of TRIM52 alone interacted with and caused degradation of TBK1, but RING domain deletion TRIM52 did not (Fig. 5B). These results suggested that TRIM52 promotes K48-linked ubiquitination of TBK1 and that the E3 ubiquitin ligase of TRIM52 is required for the ubiquitination and degradation of TBK1.

**Fig. 5.**
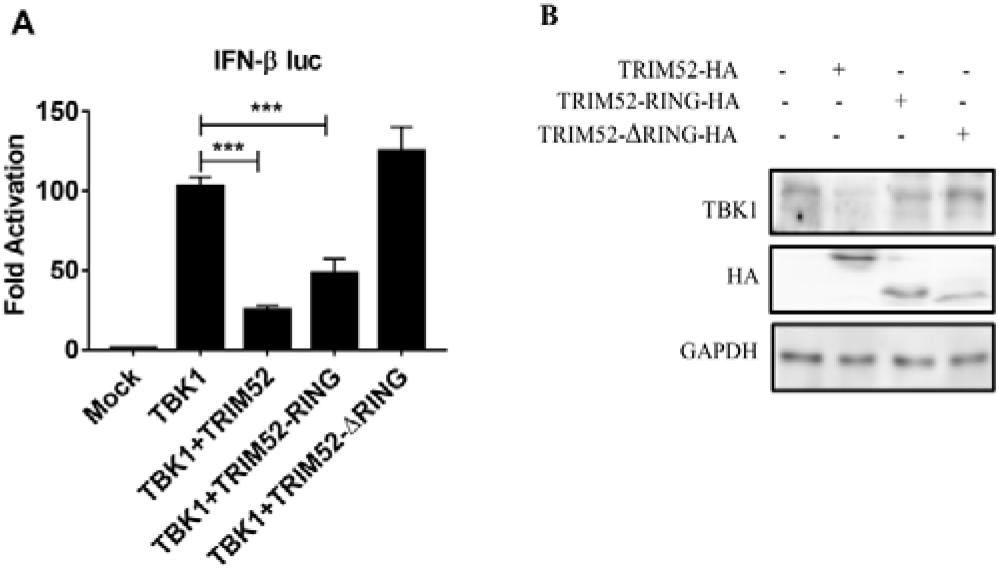
The E3 ubiquitin ligase of TRIM52 is responsible for TBK1 ubiquitination.

## DISCUSSION

Here, we demonstrated that the ubiquitin E3 ligase TRIM52 negatively regulates IFN-I signalling. Ectopic expression of TRIM52 inhibited IFN-I signalling activated by various pattern recognition receptors. It has been reported that members of the TRIM family play versatile roles in IFN-I signalling-mediated antiviral immunity [21]. Previous studies have shown that TRIM proteins such as TRIM29, TRIM30a, TRIM38, TRIM26, and TRIM39 can inhibit IFN-I production and NF-κB activation by targeting STING, NAP1, NEMO, or IRF3, respectively [15]. Hence, it is necessary to elucidate further the regulatory mechanism of innate immune signalling pathways involving the TRIM family of proteins. Our findings reveal a previously unrecognized role for TRIM52 in the inhibition of IFN-I signalling, in which TRIM52 targets TBK1 for its degradation and thus provide molecular insight into the mechanisms underlying the maintenance of innate immune homeostasis in response to viral infection.

In terms of the molecular mechanisms underlying the TRIM52-mediated inhibition of IFN-I signalling, we found that TRIM52 negatively regulated the IFN-I signalling pathway activated by all the RLR/TLR/DNA sensors, which was further supported by several lines of evidence. First, ectopic expression of TRIM52 significantly affected the production of IFN-β and ISGs induced by TBK1 but not by IRF3/5D, indicating that TRIM52 inhibited IFN-I signalling at TBK1 or upstream of TBK1. Notably, TRIM52 directly inhibits IFN-β activation by blocking IRF3 dimerization and entry into the nucleus. Second, to address how TRIM52 regulates the kinase activity of TBK1, we determined that the protein level of TRIM52 is restored by the proteasome inhibitor MG132. As an E3 ubiquitin ligase, TRIM52 has been reported to play a role in ubiquitination. In this report, we report that TRIM52 targets TBK1 for degradation through K48-linked ubiquitination by its E3 ubiquitin ligase. Research on TRIM52 has focused mainly on the regulation of tumors, and it has been reported that the expression of TRIM52 is upregulated together with that of HBx in HBV-associated HCC tissues and that TRIM52 can promote cell proliferation; moreover, the expression of TRIM52 is significantly greater in ovarian cancer, which plays an oncogenic role in ovarian cancer [22, 23]. Whether high expression of TRIM52 in tumors affects its regulation of signalling pathways remains to be further studied.

TBK1 is a key component of IFN-I signalling that is activated by various sensors of DNA and RNA. Its regulation is very important for the homeostasis of innate immune signalling, and the domain structure of TBK1 includes the kinase domain (KD), ULD, SDD, and CTD. TBK1 is activated by the phosphorylation of Ser172 in the KD domain, the dimer is stabilized by contacts between the ULD and KD with the SDD of the contralateral subunit, and the truncated CTD region contains the binding site for TANK and other adaptors [24]. In the coiled-coil domain of TBK1, Lys670 is an essential residue for NLRP4-DTX4-mediated K48-linked ubiquitination and degradation of activated TBK1 [5]. K63-linked polyubiquitination of TBK1 at K30 and K401 is required for its catalytic activation, and RNF128 promotes K63-linked ubiquitination of TBK1 at K30 and K401 [25]. Upon viral infection, TRIM9s undergo Lys-63-linked autopolyubiquitination and serve as a platform to bridge GSK3β to TBK1, leading to the activation of IRF3 signalling, and TBK1 KD is critical for binding to TRIM9s [26]. In this report, we report that TRIM52 targets TBK1 for ubiquitination and degradation, and further elucidation of the interaction region and key sites between TBK1 and TRIM52 is important.

In summary, for the first time, our findings elucidate a previously unrecognized regulation of TRIM52 targeting TBK1 for ubiquitination and degradation (Fig. 6), which may provide molecular insight into the mechanisms underlying the maintenance of innate immune homeostasis in response to viral infection and serve as a useful therapeutic target for future autoimmunity and antiviral immunity.

**Fig. 6.**
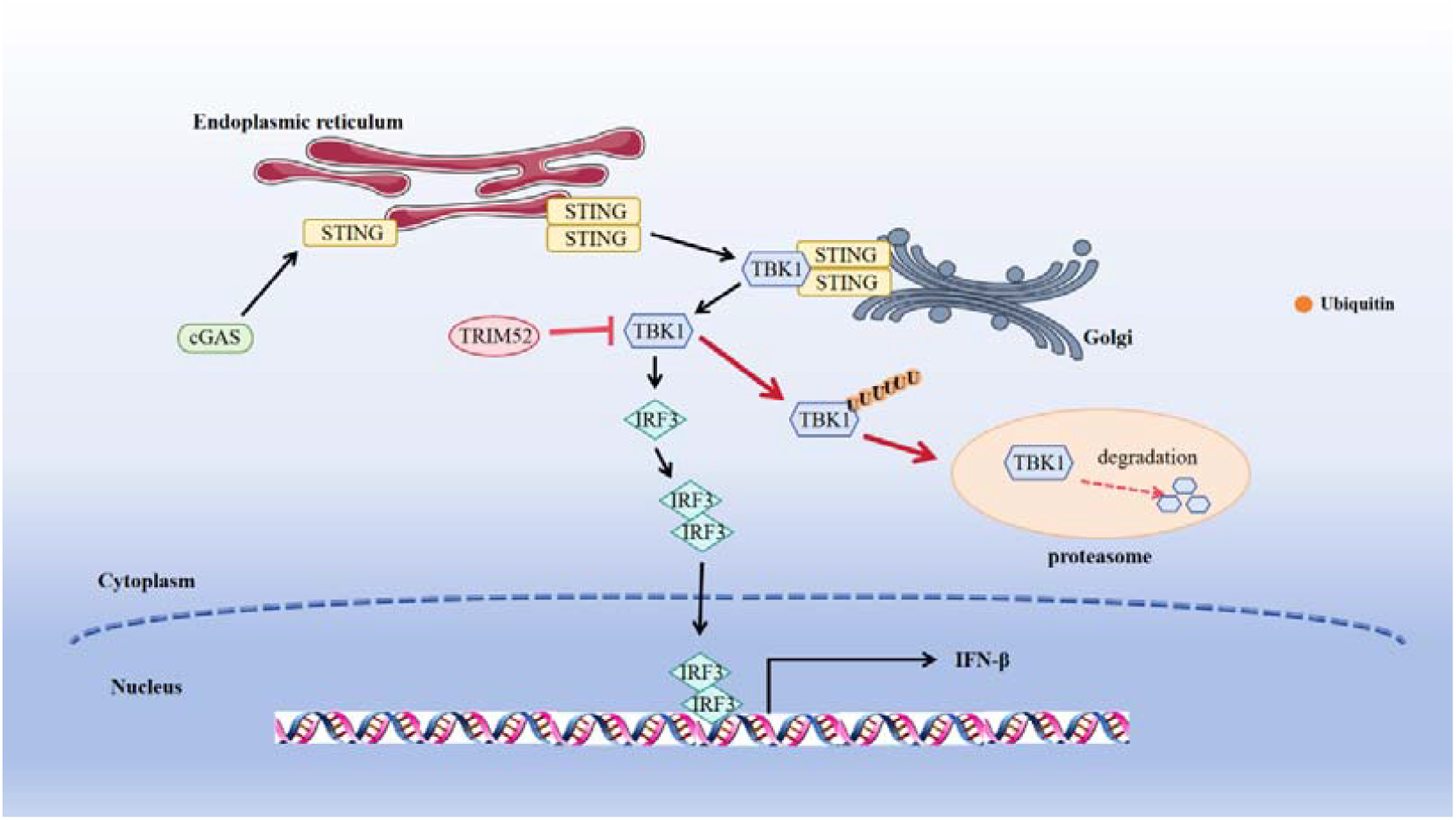
Working model of the role of TRIM52 in the cGAS-STING signalling pathway. TRIM52 targets TBK1 for ubiquitination and degradation.

## ACKNOWLEDGMENTS

We thank Rongtuan Lin for providing us with the plasmid IRF3/5D and Takashi Fujita for providing IFN-β-Luc.

